# Regulation of gene transcription by thyroid hormone receptor β agonists in clinical development for the treatment of non-alcoholic steatohepatitis (NASH)

**DOI:** 10.1101/2020.09.25.312934

**Authors:** Xuan G. Luong, Sarah K. Stevens, Andreas Jekle, Tse-I Lin, Kusum Gupta, Dinah Misner, Sushmita Chanda, Sucheta Mukherjee, Caroline Williams, Antitsa Stoycheva, Lawrence M. Blatt, Leonid N. Beigelman, Julian A. Symons, Pierre Raboisson, David McGowan, Koen Vandyck, Jerome Deval

## Abstract

Thyroid hormones are important modulators of metabolic activity in mammals and alter cholesterol and fatty acid levels through activation of the nuclear thyroid hormone receptor (THR). Currently, there are several THRβ agonists in clinical trials for the treatment of non-alcoholic steatohepatitis (NASH) that have demonstrated the potential to reduce liver fat and restore liver function. In this study, we tested three THRβ-agonism-based NASH treatment candidates, GC-1 (sobetirome), MGL-3196 (resmetirom), and VK2809, and compared their selectivity for THRβ and their ability to modulate the expression of genes specific to cholesterol and fatty acid biosynthesis and metabolism *in vitro* using human hepatic cells and *in vivo* using a rat model. Treatment with GC-1 upregulated the transcription of *CPT1A* in the human hepatocyte-derived Huh-7 cell line with a dose-response comparable to that of the native THR ligand, triiodothyronine (T3). VK2809A (active parent of VK2809), MGL-3196, and VK2809 were approximately 30-fold, 1,000-fold, and 2,000-fold less potent than T3, respectively. Additionally, these relative potencies were confirmed by quantification of other direct gene targets of THR, namely, *ANGPTL4* and *DIO1*. In primary human hepatocytes, potencies were conserved for every compound except for VK2809, which showed significantly increased potency that was comparable to that of its active counterpart, VK2809A. In high-fat diet fed rats, a single dose of T3 significantly reduced total cholesterol levels and concurrently increased liver *Dio1* and *Me1* RNA expression. MGL-3196 treatment resulted in concentration-dependent decreases in total and low-density lipoprotein cholesterol with corresponding increases in liver gene expression, but the compound was significantly less potent than T3. In conclusion, we have implemented a strategy to rank the efficacy of THRβ agonists by quantifying changes in the transcription of genes that lead to metabolic alterations, an effect that is directly downstream of THR binding and activation.

## Introduction

Non-alcoholic fatty liver disease (NAFLD), characterized by ≥ 5% hepatic fat accumulation, encompasses a heterogenous series of disorders ranging from liver steatosis to more severe non-alcoholic steatohepatitis (NASH), which may include inflammatory cell infiltration, hepatocyte ballooning, and fibrosis [1,2]. In its most severe form, NASH can progress to liver cirrhosis and hepatocellular carcinoma. Although estimates vary among studies, the worldwide prevalence of NAFLD could be as high as 25% [3]. The American Liver Foundation estimated that NAFLD is the most common cause of chronic liver disease in the United States, affecting between 80 and 100 million individuals. Twenty percent of these patients develop NASH, representing approximately 5% of total adults. Common NAFLD/NASH comorbidities include obesity, type II diabetes, hyperlipidemia, hypertension, and metabolic syndrome [3]. In the absence of any approved treatment, the medical burden and healthcare costs associated with NASH are immense.

Research on the medical treatment of NASH consists of modulating either sugar or fat metabolism or targeting one of the downstream pathways associated with liver inflammation and fibrosis [4, 5]. The largest class of molecular targets for hormone-based NASH therapies is nuclear receptors [6, 7]. There are currently several small molecule drug candidates at various stages of clinical trial evaluation. These include the farnesoid X receptor agonists, obeticholic acid and cilofexor, as well as the peroxisome proliferator-activated receptor agonists, lanifibranor, pioglitazone, elafibranor, and seladelpar. Thyroid hormone receptors (THRs) represent the third class of nuclear receptors targeted for potential NASH therapy [8, 9]. Endogenous thyroid hormones (THs), T4 and T3 (Fig 1A), are important modulators of metabolic activity in mammals and alter cholesterol and fatty acid levels through binding and activation of THRs [10]. THRs exist as two subtypes, THRα and THRβ, which are found in most tissues, but are differentially expressed [11]; THRα is highly expressed in bone and the heart, while THRβ is the major form in the liver. THRs form homodimers or heterodimerize with other nuclear receptors (e.g. retinoid X receptors, RXRs) that recognize and bind thyroid hormone response elements (TREs) located in the upstream promotor region of target genes. Upon ligand-binding, these complexes can activate or repress transcription directly, through interaction with other transcription factors, and/or *via* the recruitment of co-activators [12–14]. Here, we have characterized how THR-dependent transcription is upregulated by several thyromimetics that have reached human clinical testing for the treatment of NASH: GC-1 (sobetirome, Fig 1B), MGL-3196 (resmetirom, Fig 1C), and VK2809, a liver-targeting prodrug that is cleaved into its active parent VK2809A by cytochrome P450 isoenzyme 3A (CYP3A) after first pass intrahepatic activation (Fig 1D). GC-1 completed Phase 1 clinical trials in 2008 and demonstrated lipid-lowering effects with both single and multiple dosing [15], MGL-3196 is in Phase 3 clinical trials and has demonstrated significant reduction in hepatic fat after 12 and 36 weeks of treatment [16], and VK2809 is in Phase 2 of clinical testing and has been shown to reduce hepatic fat content in NAFLD patients after 12 weeks of treatment [17]. All three compounds have been reported to be potent and selective activators of THRβ in biochemical assays [18–20]. However, simple ligand-binding assays using truncated THR proteins do not fully recapitulate the complex THR-activation cascade that leads to changes in gene transcription and, ultimately, metabolic regulation. To this date, the characterization of THR activation by these drug candidates using *in vitro* cell-based assays that quantify gene transcription has not been reported.

**Fig 1.**
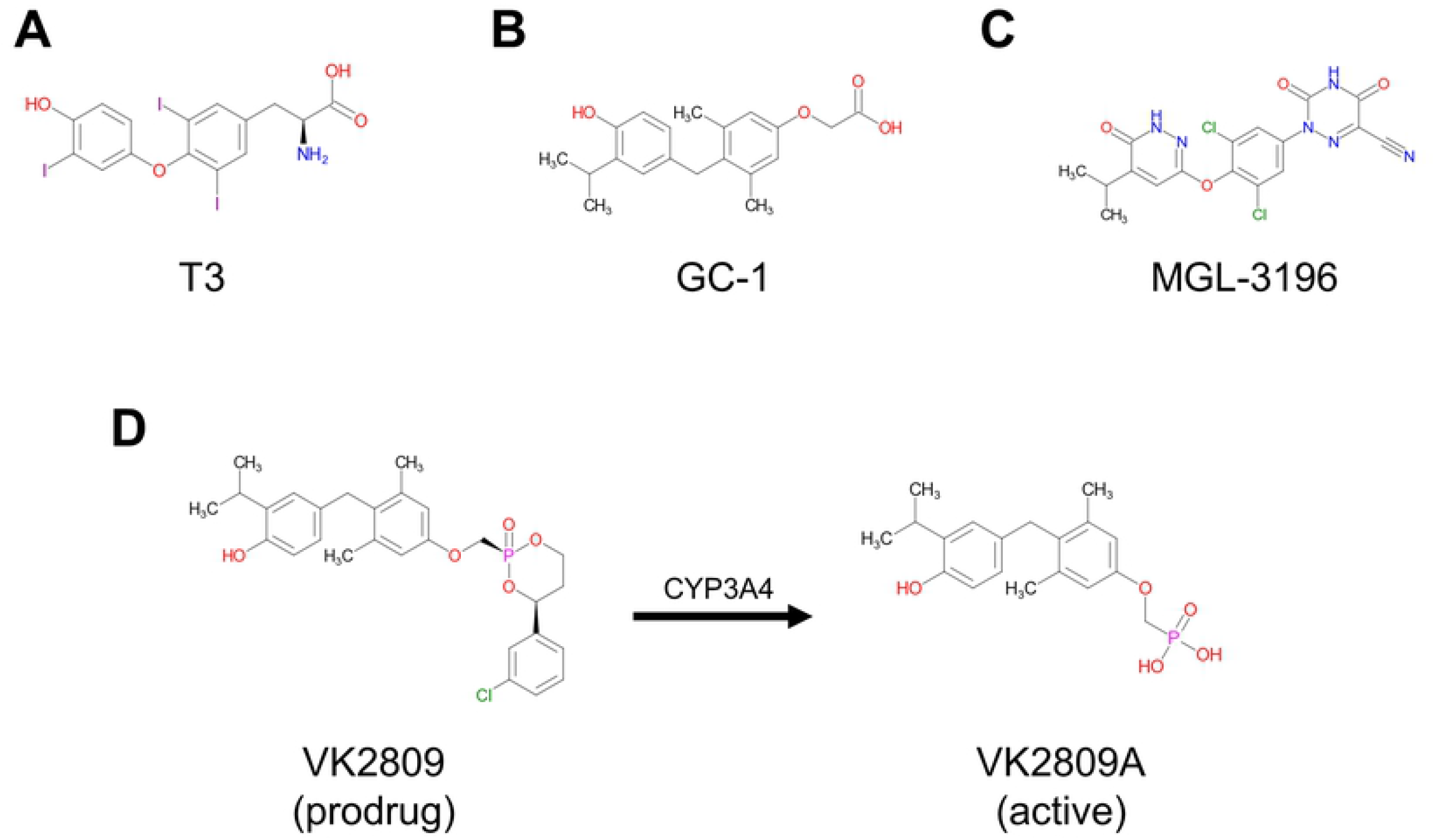
Chemical structures of test compounds. Chemical structures of (A) the natural THR ligand triiodothyronine, T3, (B) sobetirome, GC-1, (C) resmetirom, MGL-3196, (D) VK2809, and VK2809A, which is produced by CYP3A-mediated cleavage of VK2809 after first pass intrahepatic activation.

In this study, we have developed a streamlined screening cascade using *in vitro* and *in vivo* systems to evaluate the potency of thyromimetic candidates for the treatment of NASH. The aim of this study was to compare the ability of clinically relevant THRβ agonists, GC-1, MGL-3196, and VK2809, to activate their cognate receptor, modulate gene expression, and, ultimately, alter cholesterol and fatty acid biosynthesis and metabolism in the liver. We found that monitoring gene expression changes in human hepatocyte-derived cell lines and primary human hepatocytes (PHH) provides a valuable first screen of THRβ agonists and accurately predicts clinical efficacy.

## Material and methods

### Compounds

T3 (T2877) and GC-1 (SML1900) were purchased from Sigma-Aldrich. MGL-3196, VK2809, and VK2809A were synthesized by WuXi AppTec Limited (China) and compound identities and purities were verified *via* high-performance liquid chromatography and liquid chromatography–mass spectrometry. All compounds were dissolved in dimethyl sulfoxide (DMSO; Sigma-Aldrich, D4540)

### TR-FRET thyroid hormone receptor coactivator assay

A time-resolved FRET (TR-FRET)-based, biochemical assay was used as an initial screen to assess the ability of compounds to bind either THRα or β *in vitro.* Briefly, binding of an agonist to the GST-tagged THR ligand-binding domain (LBD) causes a conformational change, resulting in higher affinity for the coactivator peptide. Upon excitation of the terbium-labeled anti-GST antibody, energy is transferred to the fluorescein-labeled coactivator peptide and is detected as emission at 520 nm.

The assay procedure is based on the manufacturer’s protocol for LanthaScreen™ TR-FRET Thyroid Receptor beta Coactivator Assay (Invitrogen, PV4686) with slight, optimized modifications. Briefly, the assay was performed in 384-well, black microplate plates, protected from light. Test compounds were serially diluted in DMSO (1.0% final DMSO concentration) and added to the test plate. GST-tagged THRα or β LBD was added to the plate to yield a final concentration of 1.0 nM, followed by a mixture of the fluorescein-labeled SRC2-2 coactivator peptide and terbium-labeled anti-GST antibody at the final concentrations of 200.0 nM and 2.0 nM, respectively. After 90 mins incubation at room temperature (RT), TR-FRET was measured on a VICTOR multilabel plate reader (Perkin Elmer) using an excitation wavelength of 340 nm with 495 nm and 520 nm emission filters. The results were then quantified by expressing ratios of the intensities (520:495) and dose-response curves were fitted by non-linear regression with variable slope. Statistical analysis was performed in GraphPad Prism 8.0.

### Luciferase reporter assay in HEK293T cells

After initial characterization of *in vitro* THR-binding/activation, compounds were tested for their ability to bind and activate THRα or β (in complex with RXR), inducing gene expression, in cultured human-derived cells. To this end, HEK293T cells were transiently transfected with a firefly luciferase reporter under control of a TRE (TRE-Luc), a RXR expression plasmid, and either a THRα or β expression plasmid. This assay was performed at Pharmaron Beijing Co., Ltd. (China).

Briefly, HEK293T cells (ATCC, CRL-3216) were seeded into 6-well culture plates at 7.0 x 10^5^ cells/well and cultured in Dulbecco’s Modified Eagle’s Medium (DMEM; Hyclone, SH30022) supplemented with 10% fetal bovine serum (FBS; Gibco, 16000-044) and 1% Penicillin-Streptomycin (P/S; Corning, 30-002-CI) at 37°C and 5% CO_2_. After 24 hrs of incubation, transfection complexes were prepared by mixing 12 μL Lipofectamine 2000 (Invitrogen, 11668019) with 4 μg of a plasmid mixture (1:1:4 THR:RXR:TRE-Luc) in 200 μL Opti-MEM (Invitrogen, 11058-021) and added to the cells. After overnight incubation, the transfected cells were re-seeded at 1.0 x 10^4^ cells/well into 384-well microplates and incubated for an additional 5 to 6 hrs. Test compounds were serially diluted in DMSO and added to the cells (0.1% final DMSO concentration). After approximately 18 to 24 hrs, the culture plates were equilibrated to RT, 30 μL ONE-Glo reagent (Promega, E6120) was added to each well, and luminescence was measured on an EnSpire plate reader (Perkin Elmer). The results were then quantified by calculating percent agonism and dose-response curves were fitted by non-linear regression with variable slope. Statistical analysis was performed in GraphPad Prism 8.0.

### Differential gene expression assay in hepatic cells

Huh-7 cells (JCRB Cell Bank, JCRB0403) were routinely cultured in DMEM (Corning, 10-013-CM) supplemented with 10% FBS and 1% P/S at 37°C and 5% CO2 until 80-90% confluency. Cells were then detached with 0.05% trypsin (Corning, 25-052-CV), resuspended in TH-free medium (DMEM supplemented with 10% TH-depleted FBS and 1% P/S), and seeded into collagen-coated, 96-well microplates (Corning, 354407) at 5.0 x 10^4^ cells/well. After 24 hrs, the culture medium was replaced with treatment media. Cells were treated for 24 hrs. TH-depletion of the FBS *via* resin treatment was accomplished as previously described [21]. All treatment media were made by mixing test compounds, serially diluted in DMSO, with TH-free medium (0.1% final DMSO concentration).

Transporter certified human hepatocytes (PHH) were obtained from BioIVT (Lot: JEL, F00995-TCERT). Cells were thawed in Cryopreserved Hepatocytes Recovery Medium (Gibco, CM7000) and plated into collagen-coated, 96-well microplates at 6.0 x 10^4^ cells/well. After 6 hrs, the medium was replaced with serum-free incubation medium, William’s E Medium (Gibco, A1217601) supplemented with Primary Hepatocyte Maintenance Supplements (Gibco, CM4000). After 24 hrs, the incubation medium was replaced with treatment media. Cells were treated for 24 hrs. All treatment media were made by mixing test compounds, serially diluted in DMSO, with serum-free incubation medium (0.1% final DMSO concentration).

After 24 hrs in treatment media, both Huh-7 cells and PHH were processed with the TaqMan Fast Advanced Cells-to-Ct Kit (Invitrogen, A35378), according to the manufacturer’s protocol. Briefly, the treatment media was removed and the cells were washed with 50 μL cold 1X phosphate-buffered saline (Corning, 21-040-CM). Fifty μL lysis buffer containing DNase I was added to each well and the plate was incubated on a rotor at RT for 5 mins. Five μL stop solution was then added to each well and after another 2 mins incubation on a rotor at RT, the cell lysates were used for reverse transcription. The resulting cDNA was diluted 1:2 with nuclease-free, distilled water (Invitrogen, 10977015). Gene expression was measured using TaqMan Fast Advanced Master Mix (Applied Biosystems, 4444964) and the following TaqMan Gene Expression assays (Applied Biosystems, 4331182): *18S* (Hs99999901_s1), *ACTB* (Hs01060665_g1), *ANGPTL4* (Hs01101123_g1), *CPT1A* (Hs00912671_m1), *DIO1* (Hs00174944_m1), *TFG* (Hs02832013_g1), *THRA* (Hs00268470_m1), and *THRB* (Hs00230861_m1). *ACTB* and *TFG* served as control housekeeping genes for Huh-7 assays and *18S* and *ACTB* for PHH assays. Ten μL reactions were run on the qTOWER^3^ 84 (Analytik Jena). Relative quantification (RQ) of gene expression was calculated *via* the 2^-ΔΔCt^ method and dose-response curves were fitted by non-linear regression with variable slope. Statistical analysis was performed in GraphPad Prism 8.

### High-fat diet fed rat study

Animals were purchased from Vital River Laboratory Animal Technology Co. Ltd. and experiments were conducted at Covance Pharmaceutical R&D (Shanghai) Co., Ltd. (China). Animals were group-housed in polycarbonate cages with corncob bedding under controlled temperature (21-25°C), humidity (40-70%), and a 12-hr light/dark cycle. All procedures performed were in compliance with local animal welfare legislation, Covance global policies and procedures, and the Guide for the Care and Use of Laboratory Animals.

Male Sprague Dawley rats, approximately 8 to 11 weeks of age, were fed with either a normal diet, ND (D12450K: 10 kcal% fat, no sucrose), or high-fat diet, HFD (D12109C: 40 kcal% fat, 1.25 gm% cholesterol, 0.5 gm% sodium cholate, 12.5 gm% sucrose). After 12 days of diet consumption, baseline serum total cholesterol and low-density lipoprotein cholesterol (LDL-C) levels were measured with the cobas 6000 c501 Chemistry Analyzer (Roche) to confirm hypercholesterolemia in the HFD fed rats. Animals were then randomized into treatment groups. After 14 days, serum lipid levels were re-measured (pre-dose) and animals were orally dosed (P.O.) once with either vehicle (80% PEG400 in water) or MGL-3196 at 5.0 mg/kg, 1.5 mg/kg, or 0.5 mg/kg. T3 was administered *via* a single intraperitoneal injection (I.P.) at 0.5 mg/kg.

Twenty-four hrs after dosing, animals were euthanized by CO2 inhalation and serum and plasma were collected along with liver tissue. Serum total cholesterol and LDL-C levels were determined as described above and concentrations of compounds were measured using liquid chromatography-tandem mass spectrometry (LC-MS/MS). Tissue samples were stored in *RNAlater* (Invitrogen, AM7020) at −70°C until homogenization with the Scientz-48 TissueLyser LT. RNA extraction was performed at WuXi AppTec (Hong Kong) Limited (China) with the RNeasy Mini Kit (Qiagen, 74106) and RNA concentration and quality was determined using the Nanodrop 2000 (Thermo Scientific); additional quality control was assessed with agarose gel electrophoresis. One μg total RNA from each sample was then reverse-transcribed using the High Capacity cDNA Reverse Transcription kit (Applied Biosystems, 4368814) and the resulting cDNA was diluted 1:5 with nuclease-free, distilled water. Gene expression was measured using TaqMan Fast Advanced Master Mix and the following TaqMan Gene Expression assays: *Actb* (Rn00667869_m1), *Cpt1a* (Rn00580702_m1), *Dio1* (Rn00572183_m1), *Me1* (Rn00561502_m1), *Rplp1* (Rn03467157_gH), and *Thrsp* (Rn01511034_m1). *Actb* and *Rplp1* served as control housekeeping genes. Ten μL reactions were run on the qTOWER3 84. Relative quantification (RQ) of gene expression was calculated *via* the 2^-ΔΔCt^ method and statistical analysis was performed in GraphPad Prism 8.0.

### Ethics statement

Experiments involving rats were conducted at Covance Pharmaceutical R&D (Shanghai) Co., Ltd. All procedures performed were in compliance with local animal welfare legislation, Covance global policies and procedures, and the Guide for the Care and Use of Laboratory Animals. Animals were euthanized by CO_2_ inhalation.

## Results

### Characterization of THRα and THRβ activation

Due to the significant and broad role of THs in human development and physiology, a desirable property of NASH therapeutic thyromimetics is that their action be focused to the liver in order to decrease the risk of adverse, off-target effects on the heart, bone, and muscle [22]. This can be achieved by either targeting a compound to the liver or by increasing its selectivity for THRβ compared to THRα. Using a biochemical approach, the TR-FRET thyroid hormone receptor coactivator assay, and a cell-based approach, the HEK293T luciferase reporter assay, we characterized the ability of compounds to bind and activate each THR subtype. The EC_50_ value calculated from each dose-response curve measures the potency by which a compound activates either THR; a large THRα EC_50_-to-THRβ EC_50_ ratio (α:β) indicates increased selectively towards THRβ (Table 1).

**Table 1.** THRα/β EC_50_ values and selectivity of test compounds.

| Compound | TR-FRET assay |  |  | Luciferase reporter assay |  |  |
| --- | --- | --- | --- | --- | --- | --- |
| | THRα EC<sub>50</sub> (nM)<br>Mean ± SEM | THRβ EC<sub>50</sub> (nM)<br>Mean ± SEM | $\alpha$:$\beta$ | THRα EC<sub>50</sub> (nM)<br>Mean ± SEM | THRβ EC<sub>50</sub> (nM)<br>Mean ± SEM | $\alpha$:$\beta$ |
| **T3** | 0.3 ± 0.0<br>(n = 54) | 0.4 ± 0.0<br>(n = 54) | 0.8 | 14.3 ± 0.6<br>(n = 28) | 11.5 ± 0.6<br>(n = 27) | 1.2 |
| **GC-1** | 0.4 ± 0.1<br>(n = 3) | 0.4 ± 0.1<br>(n = 3) | 0.9 | 9.8 ± 0.6<br>(n = 29) | 4.6 ± 0.3<br>(n = 35) | 2.1 |
| **MGL-3196** | 933.8 ± 175.2<br>(n = 7) | 73.1 ± 9.3<br>(n = 7) | 12.8 | 5927.4 ± 1117.6<br>(n = 3) | 2365.8 ± 689.5<br>(n = 4) | 2.5 |
| **VK2809A** | 25.5 ± 7.0<br>(n = 3) | 10.1 ± 2.7<br>(n = 3) | 2.5 | 297.4 ± 41.4<br>(n = 4) | 269.0 ± 30.9<br>(n = 5) | 1.1 |
| **VK2809** | 253.2 ± 40.7<br>(n = 3) | 459.5 ± 163.4<br>(n = 3) | 0.6 | 54.8 ± 11.7<br>(n = 4) | 690.3 ± 28.6<br>(n = 5) | 0.1 |

In the TR-FRET thyroid hormone receptor coactivator assay, increasing concentrations of each compound was combined with the GST-tagged LBD of THRα or β, a coactivator peptide, and terbium-labeled anti-GST antibody. TR-FRET was then measured and dose-response curves were generated. In the luciferase reporter assay, HEK293T cells were transfected with a firefly luciferase reporter plasmid under the control of the TRE-Luc, a RXR expression plasmid, and either a THRα or β expression plasmid. The cells were then treated with increasing concentrations of each compound. Luminescence was measured and dose response curves were generated. For both assays, calculated EC_50_ means ± SEM, n, and THRα EC_50_-to-THRβ EC_50_ ratios (α:β) are reported for each compound-THR subtype combination.

In the TR-FRET assay, T3 showed high-affinity for both THR subtypes with no selectivity (THRα EC_50_ = 0.3 nM, THRβ EC_50_ = 0.4 nM, α:β = 0.8), as did GC-1 (THRα EC_50_ = 0.4 nM, THRβ EC_50_ = 0.4 nM, α:β = 0.9). VK2809 was a weak binder of both THRs and showed no selectivity (THRα EC_50_ = 253.2 nM, THRβ EC_50_ = 459.5 nM, α:β = 0.6), while VK2908A was relatively potent and slightly selective for THRβ (THRα EC_50_ = 25.5 nM, THRβ EC_50_ = 10.1 nM, α:β = 2.5) and MGL-3196 was the most THRβ selective compound tested, although its potency was relatively low (THRα EC_50_ = 993.8 nM, THRβ EC_50_ = 73.1 nM, α:β = 12.8).

Next, we tested whether the compounds would behave similarly in a less artificial system by indirectly measuring gene transcription changes in a cellular context with a luciferase reporter assay in HEK293T cells. In this assay, T3 remained a very potent compound, with no selectivity (THRα EC_50_ = 14.3 nM, THRβ EC_50_ = 11.5 nM, α:β = 1.2) and GC-1 was also potent with marginal THRβ selectivity (THRα EC_50_ = 9.8 nM, THRβ EC_50_ = 4.6 nM, α:β = 2.1). VK2809 showed increased potency for THRα (THRα EC_50_ = 54.8 nM, THRβ EC_50_ = 690.3 nM, α:β = 0.1), while VK2809A was considerably less potent compared to its previous characterization (THRα EC_50_ = 297.4 nM, THRβ EC_50_ = 269.0 nM, α:β = 1.1). MGL-3196 was a considerably weaker binder and activator of both THRs with reduced THRβ selectivity in this assay (THRα EC_50_ = 5927.4 nM, THRβ EC_50_ = 2365.8 nM, α:β = 2.5) compared to the TR-FRET assay.

Although both *in vitro* assays provided preliminary information on how these compounds behave, neither system is ideal. The TR-FRET assay presents a highly artificial environment using truncated versions of the THRs and the HEK293T luciferase reporter assay measures transcription indirectly *via* luciferase activity in a non-hepatocyte-derived cell-line that is overexpressing either receptor subtype.

### Differential gene expression analysis of direct THR targets in human hepatic cells

We aimed to develop an *in vitro*, cell-based assay that is not only amenable to high throughput screening of compounds, but that can also more reliably recapitulate *in vivo* processes. Because activated THR may function as a transcription factor, we compared the action of THR agonists in a human hepatocyte-derived cell line and in primary human hepatocytes (PHH) by quantifying transcriptional changes resulting from compound treatment (Fig 2A). Cells were cultured in TH-depleted media for 24 hrs and then treated with compounds for 24 hrs. The RNA levels of THR target genes were then measured *via* RT-qPCR.

**Fig 2.**
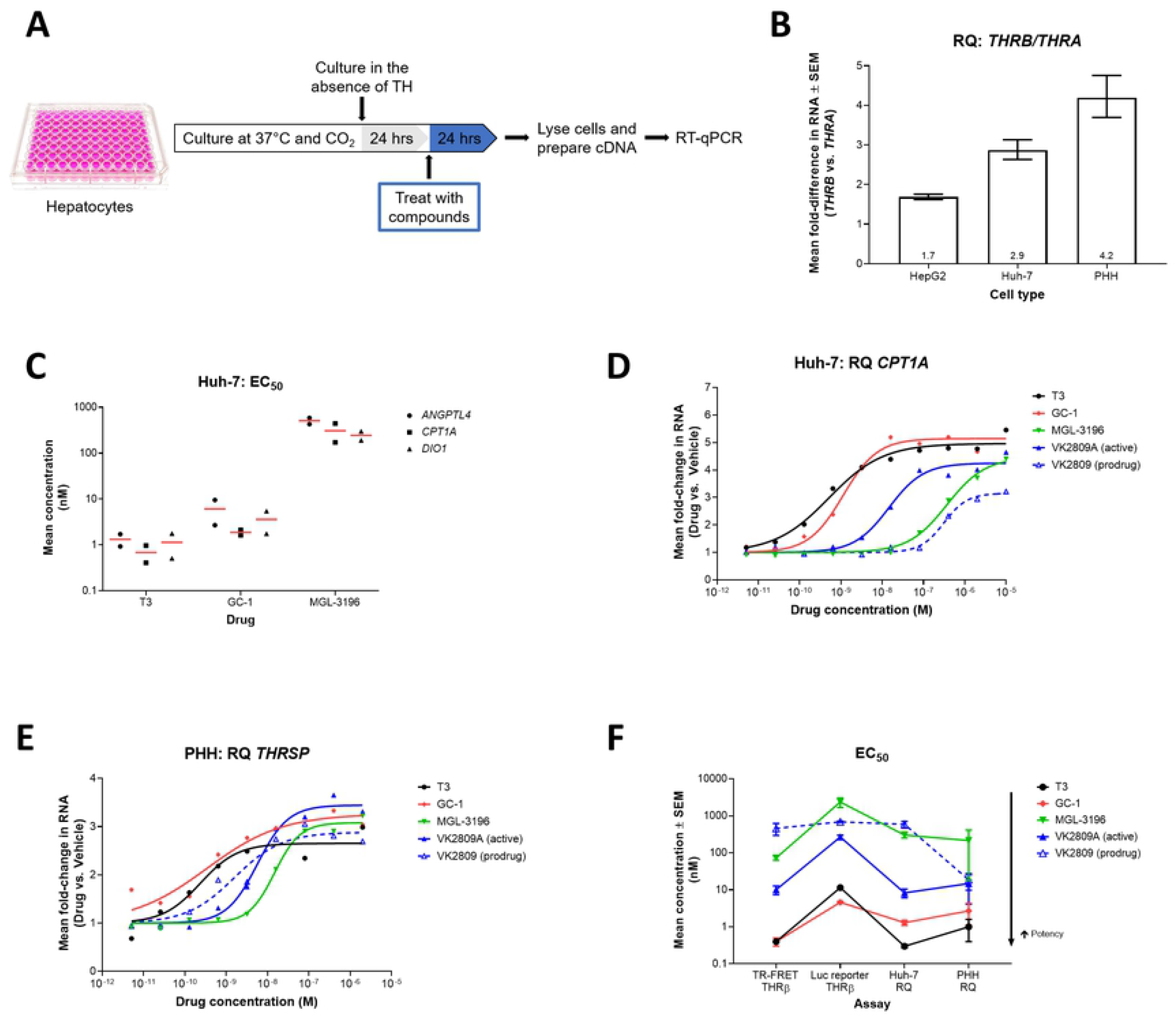
Differential gene expression in Huh-7 cells and PHH resulting from treatment with THR agonists. (A) Illustration of the *in vitro,* hepatic cell-based differential gene expression assay design. (B) *THRB* and *THRA* RNA levels were quantified by RT-qPCR in HepG2 (n = 3), Huh-7, (n = 3), and PHH (n = 5) cells. Mean RQ values ± SEM are reported with means annotated within the bars. (C) Huh-7 cells were treated with increasing doses of T3 (n = 2), GC-1 (n = 2), or MGL-3196 (n = 2) for 24 hrs. *ANGPLT4, CPT1A,* and *DIO1* RNA levels were quantified by RT-qPCR and dose-response curves were generated for each gene-compound combination. Mean EC_50_ values (red bar) and individual replicate EC_50_ values (black symbols) are reported. (D) Huh-7 cells were treated with increasing doses of T3 (black), GC-1 (red), MGL-3196 (green), VK2809A (solid blue), or VK2809 (dashed blue) for 24 hrs. *CPT1A* RNA levels were quantified by RT-qPCR. Representative mean RQ values at each compound concentration and fitted dose-response curves are reported. (E) PHH were treated with increasing doses of T3 (black), GC-1 (red), MGL-3196 (green), VK2809A (solid blue), or VK2809 (dashed blue) for 24 hrs. *THRSP* RNA levels were quantified by RT-qPCR. Representative mean RQ values at each compound concentration and fitted dose-response curves are reported. (F) EC_50_ values for every test compound were calculated from dose-response curves generated from the TR-FRET THRβ, luciferase (Luc) reporter THRβ, Huh-7 differential gene expression (RQ), and PHH RQ assays (data reported in Tables 1 and 2). Mean EC_50_ values ± SEM are reported.

To choose the most appropriate cell line for the assay, THR expression levels in HepG2 and Huh-7 cells were quantified *via* RT-qPCR. While both cell lines expressed more *THRB* than *THRA,* reflecting the expression patterns observed in liver tissue [23–26], Huh-7 cells had a larger *THRB*-to-*THRA* ratio, 2.9 compared to 1.7 for HepG2, and was thus used in downstream assays (Fig 2B). We then tested the effects of T3, GC-1, and MGL-3196 on the expression of several known THR gene targets. Treatment with these compounds resulted in dose-dependent increases in *ANGPTL4, CPT1A,* and *DIO1* transcript levels (S1A-C Figs). EC_50_ values were calculated from the resulting dose-response curves and used as a measure of potency (Fig 2C). Across the three gene targets, T3 was the most potent activator of THR (mean EC_50_ (nM): *ANGPTL4* = 1.3, *CPT1A* = 0.7, *DIO1* = 1.2), followed by GC-1 (mean EC_50_ (nM): *ANGPTL4 =* 6.2, *CPT1A* = 1.9, *DIO1* = 3.6), and MGL-3196 was the least potent compound tested (mean EC_50_ (nM): *ANGPTL4* = 508.4, *CPT1A* = 308.0, *DIO1* = 245.8). *CPT1A* transcription was ultimately chosen as the endpoint for downstream screening assays as the gene is known to be highly transcribed in the liver, a direct target of THR [27, 28], and its transcript levels were the most abundant in Huh-7 cells compared to *ANGPTL4* and *DIO1 (i.e.* lowest mean Ct value; S1D Fig). This gene encodes the enzyme, carnitine palmitoyltransferase 1A, which has an essential role in mitochondrial fatty acid β-oxidation [29, 30]. Next, we expanded testing to include other THR agonists, VK2809 and VK2809A. All compounds caused dose-dependent increases in *CPT1A* expression, but with disparate potencies (Fig 2D). EC_50_ values indicate that T3 was the most potent activator of THR, followed by GC-1, VK2809A, MGL-3196, then VK2809 (Table 2; mean Huh-7 EC_50_ (nM): 0.3, 1.3, 8.3, 303.1, and 589.1, respectively).

**Table 2.** Differential gene expression assay EC_50_ values in Huh-7 cells and PHH.

| Compound | Huh-7 EC<sub>50</sub> (nM) | PHH EC<sub>50</sub> (nM) |
| --- | --- | --- |
| | Mean ± SEM | Mean ± SEM |
| **T3** | 0.3 ± 0.03<br>(n = 22) | 1.0 ± 0.6<br>(n = 4) |
| **GC-1** | 1.3 ± 0.2<br>(n = 5) | 2.7 ± 1.6<br>(n = 4) |
| **MGL-3196** | 303.1 ± 50.9<br>(n = 17) | 216.2 ± 197.5<br>(n = 3) |
| **VK2809A** | 8.3 ± 2.2<br>(n = 5) | 14.8 ± 10.7<br>(n = 4) |
| **VK2809** | 589.1 ± 120.1<br>(n = 5) | 18.7 ± 8.9<br>(n = 3) |

Huh-7 cells and PHH were treated with increasing concentrations of each compound and resulting differential gene expression was measured and quantified as described in Figs 2D and 2E, respectively. Calculated EC_50_ means ± SEM and n are reported for each compound-cell type combination.

We next sought to validate the characterizations of these THR agonists in a more relevant cellular model, PHH. Like Huh-7 cells and liver tissue, PHH expressed 4.2-times more *THRB* than *THRA* (Fig 2B). In PHH, treatment with the THR agonists resulted in increased expression of a direct THR gene target, *THRSP* (Fig 2E), which encodes thyroid hormone responsive protein and promotes lipogenesis [31, 32]. Four out of the five compounds tested showed comparable EC_50_ values and maintained relative potencies between the two cellular models and two target genes (Fig 2F and Table 2; mean PHH EC_50_ (nM): T3 = 1.0, GC-1 = 2.7, VK2809A = 14.8, MGL-3196 = 216.2), confirming the robustness of these assays. The one exception was VK2809 (mean PHH EC_50_ = 18.7 nM), which had significantly increased potency that was almost equal to that of VK2809A in PHH.

### *In vivo* modulation of serum lipid levels and liver gene expression

We examined whether modulation of gene expression by these thyromimetics would translate into physiological alterations in metabolism. The most and least potent compounds as characterized by the *in vitro* assays, T3 and MGL-3196, respectively, were chosen to be tested *in vivo* (Fig 3A). Hypercholesterolemia was induced in rats by feeding with a HFD for two weeks. The rats were then treated with a single dose of compound and serum lipid levels and liver gene expression were quantified. *Dio1* and *Me1* are highly expressed in the liver and are known targets of THR [33–36]. DIO1, iodothyronine deiodinase 1, is a selenoprotein that functions to regulate circulating levels of T3 by catalyzing the conversion of T4 into T3 and of T3 into T2 [37, 38]. ME1, malic enzyme 1, is a NADP-dependent enzyme that generates NADPH for fatty acid biosynthesis [39, 40].

**Fig 3.**
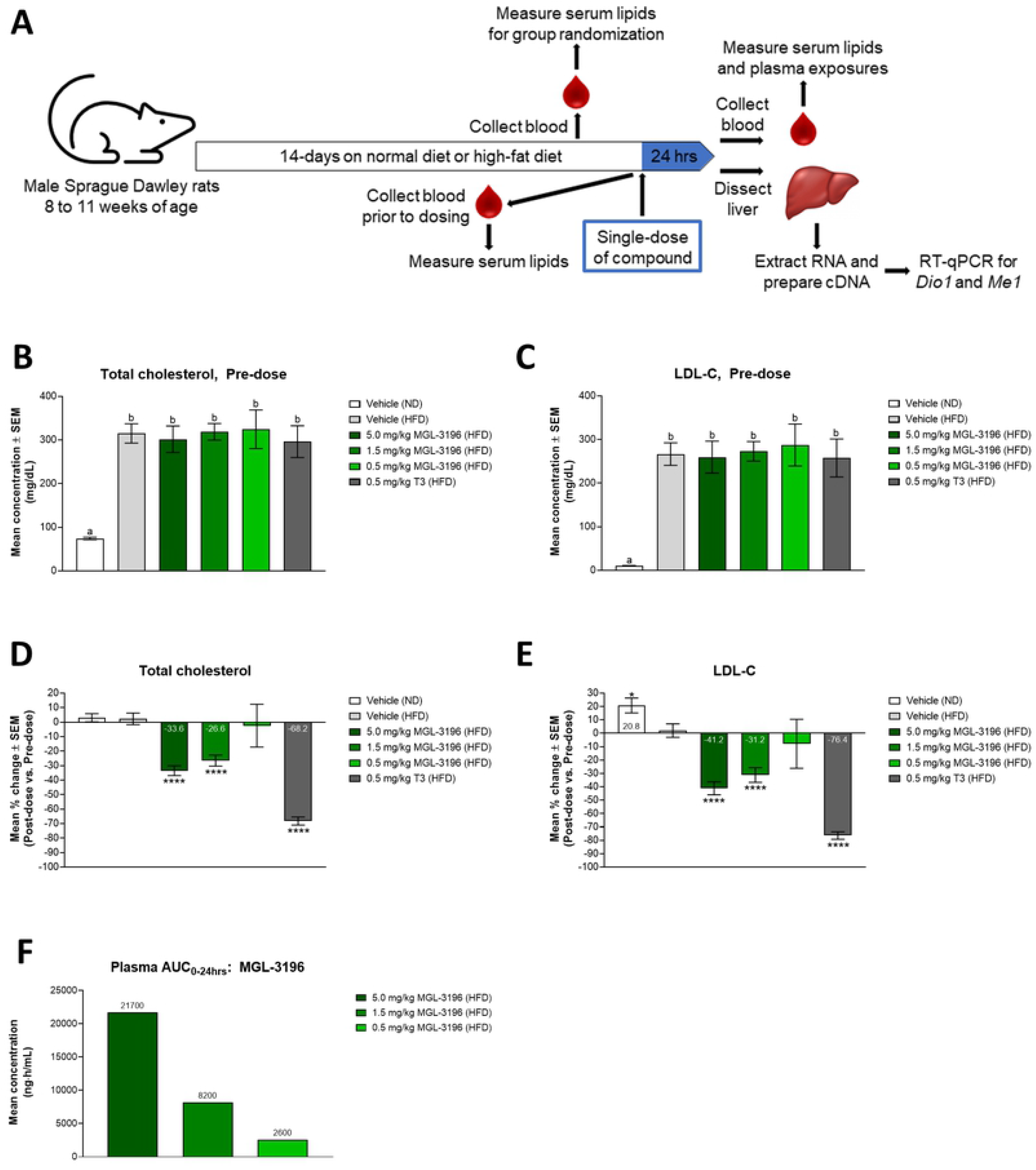
Modulation of serum lipid levels after a single-dose treatment of MGL-3196 or T3 in HFD fed rats. (A) Illustration of *in vivo* study design. (B) After two weeks of ND or HFD consumption and before compound treatment, total cholesterol levels of all animals were measured. Animals were then randomized into treatment groups: ND: vehicle (n = 20); HFD: vehicle (n = 20), 5.0 mg/kg MGL-3196 (n = 12), 1.5 mg/kg MGL-3196 (n = 20), 0.5 mg/kg MGL-3196 (n = 6), and 0.5 mg/kg T3 (n = 6). Total cholesterol level means ± SEM are reported. Statistical analysis was performed using Brown-Forsythe and Welch ANOVA and the mean of each group was compared to the mean of every other group; ‘a’ is statistically significant from ‘b’ with P < 0.01. (C) LDL-C was measured in the same animals described in B). LDL-C level means ± SEM are reported. Statistical analysis was performed using Brown-Forsythe and Welch ANOVA tests and the mean of each group was compared to the mean of every other group; ‘a’ is statistically significant from ‘b’ with P < 0.01. (D) The rats described above were then dosed once with their assigned treatments. Twenty-four hrs later, total cholesterol levels of all animals were measured. Results are presented as percent change between pre-dose and post-dose total cholesterol levels of individual animals. Percent change means ± SEM are reported with mean values annotated within the bars. Statistical analysis was performed using Brown-Forsythe and Welch ANOVA tests and the mean of each group was compared to the mean of the HFD fed, vehicle-control group; ****P < 0.0001. (E) LDL-C measurements were obtained from the same rats described in D). Results are presented as percent change between pre-dose and post-dose LDL-C levels of individual animals. Percent change means ± SEM are reported with mean values annotated within the bars. Statistical analysis was performed using Brown-Forsythe and Welch ANOVA tests and the mean of each group was compared to the mean of the HFD fed, vehicle-control group; *P < 0.05, ****P < 0.0001. (F) Plasma compound concentration in MGL-3196-treated animals was determined using LC-MS/MS. Results are presented as the mean plasma AUC_0-24hrs_ with mean values annotated above the bars.

After two weeks of feeding and prior to dosing compounds, total cholesterol and LDL-C levels were significantly elevated in HFD fed rats compared to ND fed rats, but there were no significant differences in either endpoint among the HFD groups (Figs 3B and C). Analysis comparing serum lipid levels within the same individual animals pre-dose and 24 hrs post-dose revealed that treatment with 0.5 mg/kg T3 decreased total cholesterol by 68.2%, to levels comparable to that of ND fed rats, while MGL-3196 dose-dependently decreased total cholesterol, with 5 mg/kg yielding a maximal decrease of 33.6% and 0.5 mg/kg having no significant effect (Fig 3D and S2A Fig). Similar trends were observed for LDL-C levels (Fig 3E and S2B Fig). Further analysis comparing serum lipid levels of drug-treated animals to those of HFD fed, vehicle-control animals revealed similar concentration-dependent reductions by MGL-3196 and a drastic decrease by T3 (S2C and D Figs). Moreover, pharmacokinetic analysis confirmed that exposure to MGL-3196, as measured by the area-under-the-plasma-drug-concentration-time curve (plasma AUC_0-24hrs_), was indeed linearly dose-dependent (Fig 3F).

We hypothesized that concurrent with the changes in serum lipid levels were alterations in liver gene expression induced by MGL-3196 and T3 activation of THR. Initial experiments were performed to identify reliable genetic endpoints from a panel of known THR targets, which included *Cpt1a, Dio1, Me1,* and *Thrsp.* The RNA levels of these targets were quantified in livers of HFD fed rats 4 or 24 hrs after dosing with either 5.0 or 1.5 mg/kg MGL-3196. After 4 hrs, MGL-3196 activated the transcription of *Cpt1a, Dio1,* and *Thrsp* in a dose-dependent manner, with increases in *Thrsp* levels being the most robust; there was no significant increase in *Me1* level compared to HFD fed, vehicle-control rats at this time (Fig 4A). At 24 hrs after dosing, the increases in RNA levels were dampened for *Cpt1a* and *Thrsp* (Fig 4B). By this time, however, transcription of *Dio1* and *Me1* was significantly increased with MGL-3196 treatment in a dose-dependent manner compared to HFD fed, vehicle-control rats (Fig 4B). Consequently, the expression of *Dio1* and *Me1* 24 hrs after dosing were chosen as the endpoints in subsequent experiments as these options offered the most consistent and robust responses when dosing with a thyromimetic.

**Fig 4.**
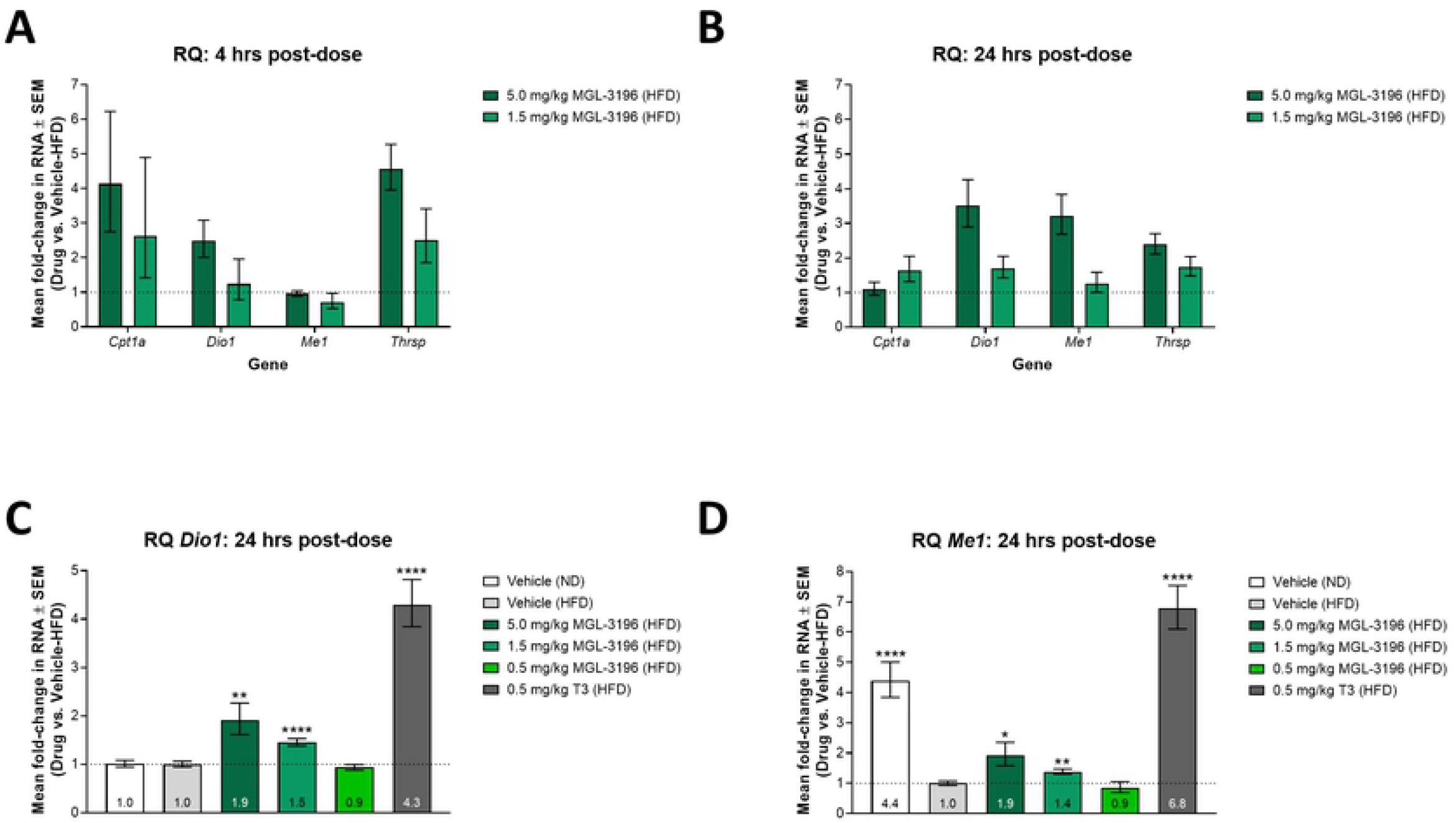
Modulation of liver gene expression after single-dose treatment of MGL-3196 or T3 in HFD fed rats. (A) After two weeks of consuming a HFD, animals were dosed once with vehicle (n = 6), 5.0 mg/kg MGL-3196 (n = 3), or 1.5 mg/kg MGL-3196 (n = 3). Four hrs later, the animals were sacrificed and liver *Cpt1a, Dio1, Me1,* and *Thrsp* RNA levels were quantified by RT-qPCR. Results are presented as expression relative to the expression levels in vehicle-control rats. Mean RQ values ± SEM are reported. (B) Animals were treated as in A) except that they were sacrificed 24 hrs post-dosing and gene expression analysis was conducted in the same manner. Results are presented as expression relative to the expression levels in vehicle-control rats. Mean RQ values ± SEM are reported. (C) The rats described in Fig 3 were sacrificed 24 hrs after compound treatment and liver *Dio1* RNA levels were quantified by RT-qPCR. Results are presented as expression relative to the expression levels in HFD fed, vehicle-control rats. Mean RQ values ± SEM are reported with mean values annotated within the bars. Statistical analysis was performed using Brown-Forsythe and Welch ANOVA tests on the ΔΔCt values. The mean of each group was compared to the mean of the HFD fed, vehicle-control group; **P < 0.01, ****P < 0.0001. (D) Liver *Me1* expression was quantified in the same rats as described in C) and in the same manner. Results are presented as expression relative to the expression levels in HFD fed, vehicle-control rats. Mean RQ values ± SEM are reported with mean values annotated within the bars. Statistical analysis was performed using Brown-Forsythe and Welch ANOVA tests on the ΔΔCt values. The mean of each group was compared to the mean of the HFD fed, vehicle-control group; *P < 0.05, **P < 0.01, ****P < 0.0001.

Liver *Dio1* and *Me1* RNA levels were quantified in the same animals whose serum lipid levels were reported in Fig 3. From these data, we were able to compare the abilities of MGL-3196 and T3 to activate THR and modulate gene transcription *in vivo.* A single dose of 0.5 mg/kg T3 resulted in a 4.3-fold increase in liver *Dio1* expression compared to the vehicle, while a single dose of MGL-3196 resulted in concentration-dependent increases in liver *Dio1* expression, with 5 mg/kg yielding the maximal 1.9 fold-increase and 0.5 mg/kg having no significant effect compared to the HFD fed, vehicle-control group (Fig 4C). Similar trends were observed for liver *Me1* expression (Fig 4D).

In summary, treatment with MGL-3196 and T3 in HFD fed rats resulted in reduced serum lipid levels that mirrored increased gene expression levels in the liver. Furthermore, we observed dose-dependent changes of these parameters with MGL-3196 treatment, which showed significantly weaker potency than T3 in the biochemical assay, cell-based assays, and in the HFD fed rat model. These results confirm that the *in vitro* characterizations of these THR agonists can be recapitulated *in vivo* and the effects of these agonists can be quantified by measuring physiological (serum lipid levels) as well as molecular (gene expression) endpoints.

## Discussion

While many studies highlight lipid level changes in animal models when profiling NAFLD/NASH therapeutic candidates, *in vitro* testing of compounds in human hepatocytes is a system that offers valuable, precursory information faster and with higher throughput. We have implemented a strategy to rank the efficacy of THRβ agonists by quantifying changes in the transcription of genes that lead to metabolic alterations, an effect that is directly downstream of THR binding and activation.

In the TR-FRET assay, GC-1 proved to be the most effective thyromimetic, with THRα/β EC_50_ values comparable to T3, followed by VK2908A, and MGL-3196 was a weak activator of either THR subtype (Table 1). The results also indicate that MGL-3196 was the most THRβ-selective thyromimetic (α:β = 12.8) and that GC-1 and VK2809A had no or minimal THRβ selectivity (α:β = 0.9 and 2.5, respectively). These findings are contrary to previously published results that describe GC-1 as having approximately 3-to 10-fold selectivity [18, 41,42] and VK2908A (MB07344) as having 15.8-fold selectivity for THRβ over THRα [43]. These discrepancies may be explained by the fact that the other studies derived selectivity from only ligand-binding affinities (*K*_d_ or *K*_i_), while the TR-FRET assay also considers coactivator recruitment, which is more physiologically relevant and a measure of receptor agonism. This type of cell-free, coactivator recruitment assay has been employed by Kelly *et al.* to characterize a variety of thyroid hormone analogues [19]. In that same study, the authors found MGL-3196 (compound 53) to be 28.3-fold more selective for THRβ over THRα, which is greater than what we observed for the compound (α:β = 12.8). This disparity could be due differences in data reporting. While we calculate selectivity as the crude THRα EC_50_-to-THR β EC_50_ ratio (α:β), Kelly *et al.* normalized this value by the selectivity of T3 from each assay. Furthermore, they reported relatively wide ranges of THR-β and THR-α values (THR-β = 0.024-0.12 μM; THR-α = 0.003-0.10 μM) compared to data reported in this current study (Table 1), which may skew their compound selectivity calculations. Other researchers using the same coactivator recruitment assay, published in a recent study data on T3, MGL-3196, and VK2809A that were consistent with our findings [44]. Kirschberg *et al*. characterized T3 and VK2809A as having no and minimal selectivity for THRβ (α:β = 1 and 2.1, respectively), while MGL-3196 had a selectivity value of 15, confirming the characterizations presented in our current study. Finally, the rankings of compound potency and THRβ selectivity as determined by the TR-FRET assay were conserved in the luciferase reporter assay for the majority of compounds tested, but all THRβ potencies were decreased in the luciferase reporter assay, especially that of MGL-3196 (Table 1 and Fig 2F).

Primary *in vitro* screens such as the TR-FRET and luciferase reporter assays provide general trends as to how test compounds may interact with THRs. However, we recognize that these approaches rely on heavily manipulated features and may not accurately reflect molecular interactions and activities occurring in hepatocytes. For example and as described above, MGL-3196 showed lower THRβ potency in the luciferase reporter assay in HEK293T cells compared to the biochemical, TR-FRET assay (32.4-fold decrease, Table 1). Furthermore, MGL-3196 was 7.8- and 10.9-times less potent in the THRβ luciferase reporter assay than in the Huh-7 and PHH gene expression assays, respectively (Fig 2F). This is due to the fact that MGL-3196 is a liver-directed drug, being a substrate for hepatic OATP1B1/B3 transporters [45]. Because these transporters are not expressed in the embryonic kidney-derived HEK293T cells, the luciferase reporter assay neglects this key feature of the compound, while the cell-free TR-FRET assay circumvents any cellular limitations. By quantifying RNA levels in human hepatocytes using RT-qPCR, we can observe gene expression changes that are directly downstream of THR-binding and activation, in a model that is more biologically relevant. However, THR subtype selectivity cannot be assessed in this more natural system.

Since employing PHH may be resource-prohibitive for many researchers, hepatocellular carcinoma (HCC) cell lines that express more *THRB* than *THRA,* such as Huh-7 cells (Fig 2B), are suitable alternatives. Furthermore, EC_50_ values from the Huh-7 differential gene expression assay were less variable than those derived from the PHH assay (Fig 2F). However, using immortalized cell lines is associated with several known caveats. In the Huh-7 assay, the prodrug VK2809 was characterized as weak activator of THR, 71-times weaker than its active parent phosphonate VK2809A (Table 2). VK2809 in PHH, however, showed drastically increased potency comparable to that of VK2809A (Fig 2F). This observation can be explained by the facts that CYP3A catalyzes the cleavage of VK2809 into VK2809A in the liver [43] and that HCC-derived cell lines have reduced CYP450 expression compared to primary hepatocytes [46]. We propose that the ineffectiveness of VK2809 in Huh-7 cells was due to the model’s decreased ability to cleave the prodrug into the active form that could then bind THR and activate gene transcription. Alternatively, in PHH, VK2809 was efficiently metabolized and, therefore, showed similar potency to VK2089A. Therefore, while HCC cell lines are an efficient system that is amenable to high-throughput screening of compounds, we recommend secondary testing of candidate thyromimetics in PHH to verify initial potency observations. These findings taken together support the use of the described Huh-7 differential gene expression assay to screen for THR agonist activity, with PHH providing increased biological relevance and additional validation. Overall, the potencies of test compounds as measured by EC_50_ were consistent across the *in vitro* assays employed in this study (Fig 2F), and the few inconsistencies were readily explained by limitations of the specific models.

We applied our preclinical screening approach to the two most advanced THR agonists in clinical development for the treatment of NASH, MGL-3196 (resmetirom), and VK2809(A). Our aim was to determine whether the behavior of MGL-3196 and VK2809(A) in our established *in vitro* and *in vivo* models were predictive of their effects in humans. To this date, the characterization of THR-mediated transcription activation by these two drug candidates using *in vitro* cell-based assays has not been reported. Gene expression upregulation by GC-1 in HepG2 cells has been previously described [23], but clinical testing of this compound for the treatment of NASH has been suspended for several years and is unlikely to resume in the near future [47]. Treatment of healthy volunteers for two weeks with MGL-3196 dosed at ≥ 80 mg resulted in 16.0-22.8% and 21.8-30.3% reduction in total and LDL-C, respectively [48]. By comparison, 14-day treatment with ≥ 5 mg of VK2809 reduced LDL-C by up to 41.2% [49]. In NASH patients, 12 weeks of treatment with MGL-3196 achieved a relative reduction in hepatic fat of approximately 36% as measured by magnetic resonance imaging-derived proton density fat fraction [16], while relative reductions in liver fat were between 53.8 and 59.7% with VK2809, depending on the dosing regimen ranging from 5-10 mg [17]. To elucidate the underlying molecular mechanisms contributing to the differences in clinical efficacy between the two molecules, we employed our described *in vitro* screening tools. In both the TR-FRET and luciferase reporter assays, the active parent phosphonate VK2809A was 7.2- to 8.8-times more potent than MGL-3196 at binding and activating THRβ (Table 1). This observation was confirmed in Huh-7 cells and in PHH, where VK2809A was 14.6- to 36.5-times more potent than MGL-3196 at activating gene transcription (Table 2). Taken together, these results confirm the ability of our *in vitro* screening methods to rapidly predict the efficacy of THR agonists in human clinical research.

Previous studies using a cholesterol-fed rat model showed that a single-dose of 0.5 mg/kg VK2809 (compound 72) resulted in a 36% reduction in total cholesterol, with up to a 55% decrease at 3.0 mg/kg [20]. In our present HFD fed rat study, a single-dose of 0.5 mg/kg of MGL-3196 did not significantly lower total or LDL-C levels (Figs 3D-E). MGL-3196 showed a dose-dependent decrease in total cholesterol starting at 1.5 mg/kg, with 5 mg/kg yielding a maximal decrease of 33.6% (Fig 3D). In comparison, a single dose of 0.5 mg/kg T3 resulted in a 68.2% decrease in total cholesterol. These results reinforce that observations that MGL-3196 has less pronounced cholesterol-lowering effects than what has been reported for VK2809. Furthermore, the cholesterol levels did not decrease linearly with MGL-3196 doses and exposures. A three-fold increase in dose, 1.5 to 5.0 mg/kg, resulted in a 2.6-fold increase in exposure (Fig 3F), but only in a 1.3-fold decrease in serum total cholesterol (Fig 3D). This plateauing effect in cholesterol reduction observed in rats is reminiscent of the lack of a linear dose-dependent cholesterol response observed in the clinic [48]. Although there could be several reasons for the plateauing dose-response of MGL-3196, the single-dose HFD fed rat model provides a valuable preclinical, *in vivo* endpoint that may predict the metabolic response in humans.

Finally, we determined that the serological effect on lipids correlates well with changes in gene expression in the liver (Fig 4). *Dio1* and *Me1* RNA levels were sensitive biomarkers of liver THR activation, with significant activation observed at 1.5 and 5.0 mg/kg MGL-3196. Again, the magnitude of MGL-3196-induced transcription was lower compared to the effect of T3, and the increase in liver *Dio1* expression between 1.5 and 5 mg/kg of MGL-3196 was only 1.3-fold (Fig 4C). Furthermore, VK2809 treatment in mice has been shown to increase the expression of CPT1A [50], a gene that we have confirmed as THR target in both Huh-7 cells and in rat liver (Fig 2D and 4A). Taken together, our *in vitro* and *in vivo* results comparing clinically relevant molecules provide a roadmap for the rapid screening of potent and selective liver targeting THRβ agonists for the potential treatment of NAFLD and NASH.

## Acknowledgments

Selected work was conducted at Covance Pharmaceutical R&D (Shanghai) Co., Ltd. by Peng Tu et al., Pharmaron Beijing Co., Ltd. by Xiaofen Guo et al., and WuXi AppTec (Hong Kong) Limited by Xin Chen, Shuang Ding, et al. We would like to thank these teams for successful collaborations.

## Supporting information captions

**S1 Fig. THR agonist dose-response curves as calculated by RQ of THR targets in Huh-7 cells**

(A) *ANGPLT4,* (B) *CPT1A,* and (C) *DIO1* RNA levels were quantified in the same cells described in Fig 2C and in the same manner. Results are presented as expression relative to the expression levels in control, vehicle-treated cells. Representative mean RQ values at each compound concentration and fitted dose-response curves are reported. (D) *ANGPLT4, CPT1A,* and *DIO1* RNA levels were quantified in the same cells described in Fig 2C and in the same manner. Ct values of each gene for control, vehicle-treated groups are reported. Results are presented as mean Ct values ± SEM.

**S2 Fig. Modulation of serum lipid levels after single-dose treatment with MGL-3196 or T3 in HFD fed rats**

(A) Total cholesterol measurements were obtained from serum of the same rats described in Fig 3D. Total cholesterol level means ± SEM are reported. Statistical analysis was performed using Brown-Forsythe and Welch ANOVA tests and the mean of each group was compared to the mean of the HFD fed, vehicle-control group; **P < 0.01, ***P < 0.001, ****P < 0.0001. (B) LDL-C measurements were obtained from serum of the same rats described in Fig 3E. LDL-C level means ± SEM are reported. Statistical analysis was performed using Brown-Forsythe and Welch ANOVA tests and the mean of each group was compared to the mean of the HFD fed, vehicle-control group; *P<0.05, **P < 0.01, ****P<0.0001. (C) Raw total cholesterol levels reported in A) were used for calculations and are the same as those used for calculations of data reported in Fig 3D. Results are presented as percent difference from the HFD fed, vehicle-control group, post-dose. Percent change means ± SEM are reported with mean values annotated within the bars. Statistical analysis was performed using Brown-Forsythe and Welch ANOVA tests and the mean of each group was compared to the mean of the HFD fed, vehicle-control group; **P<0.01,***P<0.001, ****P < 0.0001. (D) LDL-C levels reported in B) were used for calculations and are the same as those used for calculations of data reported in Fig 3E. Results are presented as percent difference from the HFD fed, vehicle-control group, post-dose. Percent change means ± SEM are reported with mean values annotated within the bars. Statistical analysis was performed using Brown-Forsythe and Welch ANOVA tests and the mean of each group was compared to the mean of the HFD fed, vehicle-control group; *P < 0.05, **P < 0.01,****P < 0.0001.

## References

1. Perumpail BJ, Khan MA, Yoo ER, Cholankeril G, Kim D, Ahmed A. Clinical epidemiology and disease burden of nonalcoholic fatty liver disease. World J Gastroenterol. 2017;23(47):8263–76. doi: 10.3748/wjg.v23.i47.8263. PubMed PMID: 29307986; PubMed Central PMCID: PMCPMC5743497.

2. Anstee QM, Targher G, Day CP. Progression of NAFLD to diabetes mellitus, cardiovascular disease or cirrhosis. Nat Rev Gastroenterol Hepatol. 2013;10(6):330–44. Epub 2013/03/19. doi: 10.1038/nrgastro.2013.41. PubMed PMID: 23507799.

3. Younossi ZM, Koenig AB, Abdelatif D, Fazel Y, Henry L, Wymer M. Global epidemiology of nonalcoholic fatty liver disease-Meta-analytic assessment of prevalence, incidence, and outcomes. Hepatology. 2016;64(1):73–84. Epub 2016/02/22. doi: 10.1002/hep.28431. PubMed PMID: 26707365.

4. Attia SL, Softic S, Mouzaki M. Evolving role for pharmacotherapy in NAFLD/NASH. Clin Transl Sci. 2020. Epub 2020/06/25. doi: 10.1111/cts.12839. PubMed PMID: 32583961.

5. Eshraghian A. Current and emerging pharmacological therapy for non-alcoholic fatty liver disease. World J Gastroenterol. 2017;23(42):7495–504. doi: 10.3748/wjg.v23.i42.7495. PubMed PMID: 29204050; PubMed Central PMCID: PMCPMC5698243.

6. Musso G, Cassader M, Gambino R. Non-alcoholic steatohepatitis: emerging molecular targets and therapeutic strategies. Nat Rev Drug Discov. 2016;15(4):249–74. Epub 2016/01/22. doi: 10.1038/nrd.2015.3. PubMed PMID: 26794269.

7. Kim KH, Lee MS. Pathogenesis of Nonalcoholic Steatohepatitis and Hormone-Based Therapeutic Approaches. Front Endocrinol (Lausanne). 2018;9:485. Epub 2018/08/24. doi: 10.3389/fendo.2018.00485. PubMed PMID: 30197624; PubMed Central PMCID: PMCPMC6117414.

8. Jakobsson T, Vedin LL, Parini P. Potential Role of Thyroid Receptor β Agonists in the Treatment of Hyperlipidemia. Drugs. 2017;77(15):1613–21. doi: 10.1007/s40265-017-0791-4. PubMed PMID: 28865063; PubMed Central PMCID: PMCPMC5613055.

9. Saponaro F, Sestito S, Runfola M, Rapposelli S, Chiellini G. Selective Thyroid Hormone Receptor-Beta (TRβ) Agonists: New Perspectives for the Treatment of Metabolic and Neurodegenerative Disorders. Front Med (Lausanne). 2020;7:331. Epub 2020/07/09. doi: 10.3389/fmed.2020.00331. PubMed PMID: 32733906; PubMed Central PMCID: PMCPMC7363807.

10. Sinha RA, Singh BK, Yen PM. Direct effects of thyroid hormones on hepatic lipid metabolism. Nat Rev Endocrinol. 2018;14(5):259–69. Epub 2018/02/23. doi: 10.1038/nrendo.2018.10. PubMed PMID: 29472712; PubMed Central PMCID: PMCPMC6013028.

11. Brent GA. Mechanisms of thyroid hormone action. J Clin Invest. 2012;122(9):3035–43. Epub 2012/09/04. doi: 10.1172/JCI60047. PubMed PMID: 22945636; PubMed Central PMCID: PMCPMC3433956.

12. Oetting A, Yen PM. New insights into thyroid hormone action. Best Pract Res Clin Endocrinol Metab. 2007;21(2):193–208. doi: 10.1016/j.beem.2007.04.004. PubMed PMID: 17574003.

13. Chan IH, Privalsky ML. Isoform-specific transcriptional activity of overlapping target genes that respond to thyroid hormone receptors alpha1 and beta1. Mol Endocrinol. 2009;23(11):1758–75. Epub 2009/07/23. doi: 10.1210/me.2009-0025. PubMed PMID: 19628582; PubMed Central PMCID: PMCPMC2775939.

14. Mengeling BJ, Lee S, Privalsky ML. Coactivator recruitment is enhanced by thyroid hormone receptor trimers. Mol Cell Endocrinol. 2008;280(1-2):47–62. Epub 2007/10/06. doi: 10.1016/j.mce.2007.09.011. PubMed PMID: 18006144; PubMed Central PMCID: PMCPMC2197157.

15. Scanlan TS. Sobetirome: a case history of bench-to-clinic drug discovery and development. Heart Fail Rev. 2010;15(2):177–82. Epub 2008/11/11. doi: 10.1007/s10741-008-9122-x. PubMed PMID: 19002578.

16. Harrison SA, Bashir MR, Guy CD, Zhou R, Moylan CA, Frias JP, et al. Resmetirom (MGL-3196) for the treatment of non-alcoholic steatohepatitis: a multicentre, randomised, double-blind, placebo-controlled, phase 2 trial. Lancet. 2019;394(10213):2012–24. Epub 2019/11/11. doi: 10.1016/S0140-6736(19)32517-6. PubMed PMID: 31727409.

17. Loomba R, Neutel J, Mohseni R, Bernard D, Severance R, Dao M, et al. LBP-20-VK2809, a Novel Liver-Directed Thyroid Receptor Beta Agonist, Significantly Reduces Liver Fat with Both Low and High Doses in Patients with Non-Alcoholic Fatty Liver Disease: A Phase 2 Randomized, Placebo-Controlled Trial. Journal of Hepatology. 2019;70(1, Supplement):e150–e1. doi: https://doi.org/10.1016/S0618-8278(19)30266-X.

18. Chiellini G, Apriletti JW, Yoshihara HA, Baxter JD, Ribeiro RC, Scanlan TS. A high-affinity subtype-selective agonist ligand for the thyroid hormone receptor. Chem Biol. 1998;5(6):299–306. doi: 10.1016/s1074-5521(98)90168-5. PubMed PMID: 9653548.

19. Kelly MJ, Pietranico-Cole S, Larigan JD, Haynes NE, Reynolds CH, Scott N, et al. Discovery of 2-[3,5-dichloro-4-(5-isopropyl-6-oxo-1,6-dihydropyridazin-3-yloxy)phenyl]-3,5-dioxo-2,3,4,5-tetrahydro[1,2,4]triazine-6-carbonitrile (MGL-3196), a Highly Selective Thyroid Hormone Receptor β agonist in clinical trials for the treatment of dyslipidemia. J Med Chem. 2014;57(10):3912–23. Epub 2014/04/08. doi: 10.1021/jm4019299. PubMed PMID: 24712661.

20. Boyer SH, Jiang H, Jacintho JD, Reddy MV, Li H, Li W, et al. Synthesis and biological evaluation of a series of liver-selective phosphonic acid thyroid hormone receptor agonists and their prodrugs. J Med Chem. 2008;51(22):7075–93. doi: 10.1021/jm800824d. PubMed PMID: 18975928.

21. Samuels HH, Stanley F, Casanova J. Depletion of L-3,5,3’-triiodothyronine and L-thyroxine in euthyroid calf serum for use in cell culture studies of the action of thyroid hormone. Endocrinology. 1979;105(1):80–5. doi: 10.1210/endo-105-1-80. PubMed PMID: 446419.

22. Pedrelli M, Pramfalk C, Parini P. Thyroid hormones and thyroid hormone receptors: effects of thyromimetics on reverse cholesterol transport. World J Gastroenterol. 2010;16(47):5958–64. doi: 10.3748/wjg.v16.i47.5958. PubMed PMID: 21157972; PubMed Central PMCID: PMCPMC3007105.

23. Yuan C, Lin JZ, Sieglaff DH, Ayers SD, Denoto-Reynolds F, Baxter JD, et al. Identical gene regulation patterns of T3 and selective thyroid hormone receptor modulator GC-1. Endocrinology. 2012;153(1):501–11. Epub 2011/11/08. doi: 10.1210/en.2011-1325. PubMed PMID: 22067320; PubMed Central PMCID: PMCPMC3249679.

24. Cheng SY, Leonard JL, Davis PJ. Molecular aspects of thyroid hormone actions. Endocr Rev. 2010;31(2):139–70. Epub 2010/01/05. doi: 10.1210/er.2009-0007. PubMed PMID: 20051527; PubMed Central PMCID: PMCPMC2852208.

25. Cheng SY. Multiple mechanisms for regulation of the transcriptional activity of thyroid hormone receptors. Rev Endocr Metab Disord. 2000;1(1-2):9–18. doi: 10.1023/a:1010052101214. PubMed PMID: 11704997.

26. Williams GR. Cloning and characterization of two novel thyroid hormone receptor beta isoforms. Mol Cell Biol. 2000;20(22):8329–42. doi: 10.1128/mcb.20.22.8329-8342.2000. PubMed PMID: 11046130; PubMed Central PMCID: PMCPMC102140.

27. Jansen MS, Cook GA, Song S, Park EA. Thyroid hormone regulates carnitine palmitoyltransferase Ialpha gene expression through elements in the promoter and first intron. J Biol Chem. 2000;275(45):34989–97. doi: 10.1074/jbc.M001752200. PubMed PMID: 10956641.

28. Thakran S, Sharma P, Attia RR, Hori RT, Deng X, Elam MB, et al. Role of sirtuin 1 in the regulation of hepatic gene expression by thyroid hormone. J Biol Chem. 2013;288(2):807–18. Epub 2012/12/03. doi: 10.1074/jbc.M112.437970. PubMed PMID: 23209300; PubMed Central PMCID: PMCPMC3543030.

29. McGarry JD, Brown NF. The mitochondrial carnitine palmitoyltransferase system. From concept to molecular analysis. Eur J Biochem. 1997;244(1):1–14. doi: 10.1111/j.1432-1033.1997.00001.x. PubMed PMID: 9063439.

30. Lee K, Kerner J, Hoppel CL. Mitochondrial carnitine palmitoyltransferase 1a (CPT1a) is part of an outer membrane fatty acid transfer complex. J Biol Chem. 2011;286(29):25655–62. Epub 2011/05/26. doi: 10.1074/jbc.M111.228692. PubMed PMID: 21622568; PubMed Central PMCID: PMCPMC3138250.

31. Wu J, Wang C, Li S, Wang W, Li J, Chi Y, et al. Thyroid hormone-responsive SPOT 14 homolog promotes hepatic lipogenesis, and its expression is regulated by liver X receptor α through a sterol regulatory element-binding protein 1c-dependent mechanism in mice. Hepatology. 2013;58(2):617–28. Epub 2013/07/02. doi: 10.1002/hep.26272. PubMed PMID: 23348573.

32. Anderson GW, Zhu Q, Metkowski J, Stack MJ, Gopinath S, Mariash CN. The Thrsp null mouse (Thrsp(tm1cnm)) and diet-induced obesity. Mol Cell Endocrinol. 2009;302(1):99–107. Epub 2009/01/20. doi: 10.1016/j.mce.2009.01.005. PubMed PMID: 19356628; PubMed Central PMCID: PMCPMC2671690.

33. Maia AL, Kieffer JD, Harney JW, Larsen PR. Effect of 3,5,3’-Triiodothyronine (T3) administration on dio1 gene expression and T3 metabolism in normal and type 1 deiodinase-deficient mice. Endocrinology. 1995;136(11):4842–9. doi: 10.1210/endo.136.11.7588215. PubMed PMID: 7588215.

34. Zavacki AM, Ying H, Christoffolete MA, Aerts G, So E, Harney JW, et al. Type 1 iodothyronine deiodinase is a sensitive marker of peripheral thyroid status in the mouse. Endocrinology. 2005;146(3):1568–75. Epub 2004/12/09. doi: 10.1210/en.2004-1392. PubMed PMID: 15591136.

35. Goodridge AG, Adelman TG. Regulation of malic enzyme synthesis by insulin triiodothyronine, and glucagon in liver cells in culture. J Biol Chem. 1976;251(10):3027–32. PubMed PMID: 944696.

36. Towle HC, Mariash CN, Oppenheimer JH. Changes in the hepatic levels of messenger ribonucleic acid for malic enzyme during induction by thyroid hormone or diet. Biochemistry. 1980;19(3):579–85. doi: 10.1021/bi00544a029. PubMed PMID: 7356948.

37. Bianco AC, Salvatore D, Gereben B, Berry MJ, Larsen PR. Biochemistry, cellular and molecular biology, and physiological roles of the iodothyronine selenodeiodinases. Endocr Rev. 2002;23(1):38–89. doi: 10.1210/edrv.23.1.0455. PubMed PMID: 11844744.

38. Schneider MJ, Fiering SN, Thai B, Wu SY, St Germain E, Parlow AF, et al. Targeted disruption of the type 1 selenodeiodinase gene (Dio1) results in marked changes in thyroid hormone economy in mice. Endocrinology. 2006;147(1):580–9. Epub 2005/10/13. doi: 10.1210/en.2005-0739. PubMed PMID: 16223863.

39. Iritani N. Nutritional and hormonal regulation of lipogenic-enzyme gene expression in rat liver. Eur J Biochem. 1992;205(2):433–42. doi: 10.1111/j.1432-1033.1992.tb16797.x. PubMed PMID: 1349281.

40. Shimomura I, Shimano H, Korn BS, Bashmakov Y, Horton JD. Nuclear sterol regulatory element-binding proteins activate genes responsible for the entire program of unsaturated fatty acid biosynthesis in transgenic mouse liver. J Biol Chem. 1998;273(52):35299–306. doi: 10.1074/jbc.273.52.35299. PubMed PMID: 9857071.

41. Wagner RL, Huber BR, Shiau AK, Kelly A, Cunha Lima ST, Scanlan TS, et al. Hormone selectivity in thyroid hormone receptors. Mol Endocrinol. 2001;15(3):398–410. doi: 10.1210/mend.15.3.0608. PubMed PMID: 11222741.

42. Haning H, Woltering M, Mueller U, Schmidt G, Schmeck C, Voehringer V, et al. Novel heterocyclic thyromimetics. Bioorg Med Chem Lett. 2005;15(7):1835–40. doi: 10.1016/j.bmcl.2005.02.028. PubMed PMID: 15780617.

43. Erion MD, Cable EE, Ito BR, Jiang H, Fujitaki JM, Finn PD, et al. Targeting thyroid hormone receptor-beta agonists to the liver reduces cholesterol and triglycerides and improves the therapeutic index. Proc Natl Acad Sci U S A. 2007;104(39):15490–5. Epub 2007/09/18. doi: 10.1073/pnas.0702759104. PubMed PMID: 17878314; PubMed Central PMCID: PMCPMC1978486.

44. Kirschberg TA, Jones CT, Xu Y, Fenaux M, Halcomb RL, Wang Y, et al. Selective Thyroid Hormone Receptor β Agonists with Oxadiazolone Acid Isosteres. Bioorganic & Medicinal Chemistry Letters. 2020:127465. doi: https://doi.org/10.1016/j.bmcl.2020.127465.

45. Kalliokoski A, Niemi M. Impact of OATP transporters on pharmacokinetics. Br J Pharmacol. 2009;158(3):693–705. Epub 2009/09/25. doi: 10.1111/j.1476-5381.2009.00430.x. PubMed PMID: 19785645; PubMed Central PMCID: PMCPMC2765590.

46. Rodríguez-Antona C, Donato MT, Boobis A, Edwards RJ, Watts PS, Castell JV, et al. Cytochrome P450 expression in human hepatocytes and hepatoma cell lines: molecular mechanisms that determine lower expression in cultured cells. Xenobiotica. 2002;32(6):505–20. doi: 10.1080/00498250210128675. PubMed PMID: 12160483.

47. Columbano A, Chiellini G, Kowalik MA. GC-1: A Thyromimetic With Multiple Therapeutic Applications in Liver Disease. Gene Expr. 2017;17(4):265–75. Epub 2017/06/13. doi: 10.3727/105221617X14968563796227. PubMed PMID: 28635586; PubMed Central PMCID: PMCPMC5885148.

48. Taub R, Chiang E, Chabot-Blanchet M, Kelly MJ, Reeves RA, Guertin MC, et al. Lipid lowering in healthy volunteers treated with multiple doses of MGL-3196, a liver-targeted thyroid hormone receptor-β agonist. Atherosclerosis. 2013;230(2):373–80. Epub 2013/08/21. doi: 10.1016/j.atherosclerosis.2013.07.056. PubMed PMID: 24075770.

49. Lian B, Hanley R, Schoenfeld S. A PHASE 1 RANDOMIZED, DOUBLE-BLIND, PLACEBO-CONTROLLED, MULTIPLE ASCENDING DOSE STUDY TO EVALUATE SAFETY, TOLERABILITY AND PHARMACOKINETICS OF THE LIVER-SELECTIVE TR-BETA AGONIST VK2809 (MB07811) IN HYPERCHOLESTEROLEMIC SUBJECTS. Journal of the American College of Cardiology. 2016;67(13 Supplement):1932. doi: 10.1016/s0735-1097(16)31933-7.

50. Zhou J, Waskowicz LR, Lim A, Liao XH, Lian B, Masamune H, et al. A Liver-Specific Thyromimetic, VK2809, Decreases Hepatosteatosis in Glycogen Storage Disease Type Ia. Thyroid. 2019;29(8):1158–67. doi: 10.1089/thy.2019.0007. PubMed PMID: 31337282; PubMed Central PMCID: PMCPMC6707038.

